# Evolutionary conservation and divergence of the transcriptional regulation of bivalve shell secretion across life history stages

**DOI:** 10.1101/2022.04.22.489168

**Authors:** Alessandro Cavallo, Melody S. Clark, Lloyd S. Peck, Elizabeth M. Harper, Victoria A. Sleight

## Abstract

Adult molluscs produce shells with diverse morphologies and ornamentations, different colour patterns and microstructures. The larval shell however, is a phenotypically more conserved structure. How do developmental and evolutionary processes generate varying diversity at different life history stages? Using live-imaging, histology, scanning electron microscopy and transcriptomic profiling, we have described shell development in a heteroconchian bivalve the Antarctic clam, *Laternula elliptica* and compared it to adult shell secretion processes in the same species. Adult downstream shell genes, such as those encoding extracellular matrix proteins and biomineralisation enzymes, were largely not expressed during shell development, and instead, a development-specific downstream gene repertoire was expressed. Upstream regulatory genes such as transcription factors and signalling molecules were conserved between developmental and adult shell secretion. Comparing heteroconchian transcriptomic data with recently reported pteriomorphian larval shell proteome data suggests that, despite being phenotypically more conserved, the downstream effectors constituting the larval shell “tool-kit” may be as diverse as that of adults. Overall, our new data suggests that a larval shell formed using development-specific downstream effector genes is a conserved and ancestral feature of the bivalve lineage, and possibly more broadly across the molluscs.

## Introduction

Molluscan shells are environmentally, economically and evolutionarily important (McDougall and Degnan 2018). The hard multifunctional external shells of molluscs are often attributed to the evolutionary success of this group. Shells are exquisitely preserved in the fossil record and their expansive extant adaptive diversity of form, as well as remarkable phenotypic plasticity, provides a powerful system to study morphological evolution (Thompson 1992). In the last decade, deciphering the molecular mechanisms that control shell secretion has received particular focus (Clark et al. 2020). A range of gastropod (Herlitze et al. 2018; Marie et al. 2010), cephalopod (Marie et al. 2009; Setiamarga et al. 2020) and bivalve (Arivalagan et al. 2017; Sleight et al. 2016a) shells have been subject to proteomic sequencing, coupled to the transcriptomic investigation of the shell-secreting mantle tissue (Arivalagan et al. 2016; Sleight et al. 2016b), resulting in large lists of candidate shell-forming genes and proteins. Comparative methods have found that despite the “deep” homology of molluscan shells and shell plates (Vinther 2015), the molecular mechanisms that control biomineralisation in molluscs – particularly the downstream effectors such as the shell matrix proteins - are extraordinarily diverse (Jackson et al. 2006). Features such as repeat low complexity domains (RLCD), domain shuffling and co-option (Aguilera et al. 2017), gene family expansions and subsequent subfunctionalisation (Aguilera et al. 2014) drive much of the observed molecular diversity. A handful of core protein domains however, are essential for all molluscan biomineralisation, regardless of morphologies, microstructures or polymorphs (carbonic anhydrase, chitin-binding, VWA and tyrosinase domains; Arivalagan et al. 2017). Due to the likelihood of fossilization, ease of study and ability to collect sufficiently large samples, there is an overwhelming bias towards adult shell studies, particularly in proteomic and transcriptomic studies. In developmental studies, some candidate gene approaches have been successfully used (Hohagen and Jackson 2013; Liu et al. 2017; 2020; Nederbragt et al. 2002), but comparatively little is known about the molecular processes that pattern and generate the embryonic shell field, larval mantle and early shell secretion.

Phenotypically, larval mollusc shells are more conserved than adult shells in terms of morphology, crystal polymorph and microstructure; they are unsculptured simplified forms composed of an organic outer layer and aragonite mineralised layer (Weiss et al. 2002). More generally, developmental processes are highly conserved and under strong selective constraints (Prud’homme and Gompel 2010; Smith et al. 1985) and so, if there are deeply homologous molecular mechanisms directing the production of the molluscan shell, they are more likely present during developmental stages. Although limited in number and taxonomic coverage, studies on the molecular control of shell development have uncovered intriguing insights. Recent work in pteriomorphian bivalves (oysters and mussels) compared the proteomes of larval shells to that of adults in the same species and showed that the repertoire of extracellular matrix proteins in the larval shell is almost completely different to that of the adult shell (Carini et al. 2019; Zhao et al. 2018). Only the handful of core protein domains that have been previously described as the core “tool-kit” for molluscan biomineralisation were shared between the two life history stages (carbonic anhydrase, chitin-binding and VWA). To date, intraspecific comparisons of the molecular mechanisms governing larval versus adult shell production are restricted to one bivalve infraclass (Pteriomorphia, e.g. oysters and mussels). These studies exclusively used proteomics in a qualitative presence vs absence approach, with non-replicated estimates of transcript abundance through ontogeny. They were the first studies to reveal the striking difference between larval and adult shells at the molecular level and they focus solely on the post-translationally modified downstream effectors, ie the shell matrix proteins. Much less is known about the transcriptional mechanisms that regulate early shell development, in comparison with the adult shell of the same species.

Here we studied shell development using imaging and quantitative transcriptomics in a member of the Heteroconchia infraclass, the Antarctic clam, *Laternula elliptica*. Heteroconchia split from the rest of the Autobranchia in the Cambrian Period, around 500 million years ago (mya; Plazzi and Passamonti 2010), and so comparisons within the Autobranchia but between Heteroconchia and Pteriomorphia shed light on features that are deeply conserved within the bivalves over 500 million years of evolutionary time.

## Results

### Morphological mantle and shell landmarks through development in *Laternula elliptica*

To phenotypically characterise the development of the shell field, larval mantle and shell in *L. elliptica*, a combination of live light imaging, histological staining and scanning electron microscopy was used (Fig. 1). Phenotypic characterisations allowed us to assign key mantle and shell landmarks to each of the development stages studied (mantle fold appearance, organic vs mineralised shell, prodissoconch I vs prodissoconch II vs dissoconch I shell) to use as a framework to decipher the transcriptional regulation of shell secretion.

**Figure 1.**
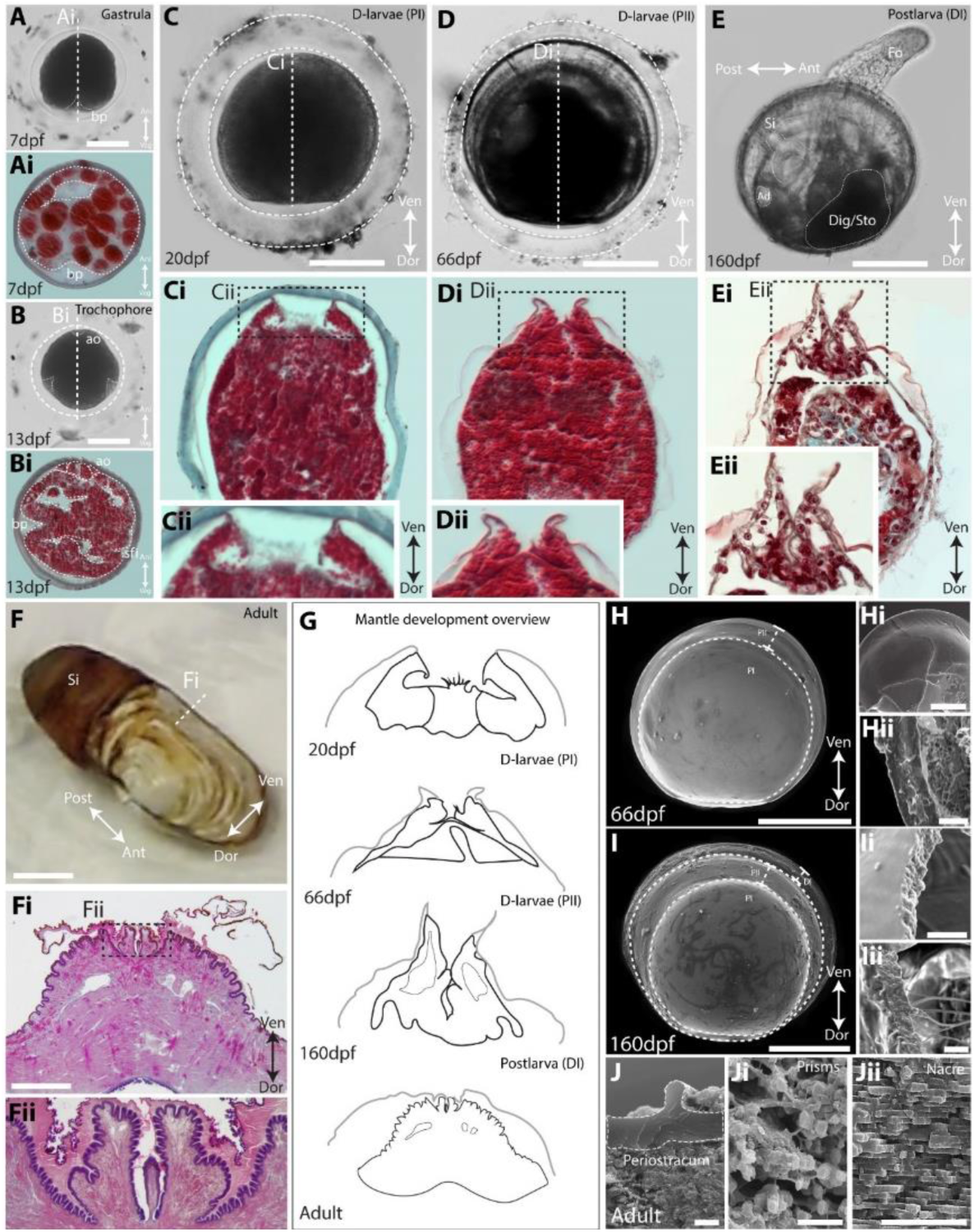
Shell and mantle development in *Laternula elliptica* characterised using live imaging, histology and Scanning Electron Microscopy (SEM). **A-Ai** Invagination of the blastopore (bp) during gastrulation 7 days post fertilization (dpf) forming the archenteron prior to shell field induction (encapsulated, scale bar 60µm). **B-Bi** Early trochophore 13dpf apical organ (ao) and shell field invagination (sfi) (encapsulated, scale bar 60µm). **C-Cii** Early D-stage larvae 20dpf, first prodissoconch I organic shell (PI) secreted by unfolded larval mantle (Ci-Cii), reduced ciliary velum resorbing (Cii) (encapsulated, scale bar 60µm). **D-Dii** Late D-stage larvae 66dpf with prodissoconch II (PII) that is mineralized, mantle folds appear and fuse to form the early fused inner mantle fold (capsule disintegrating, scale bar 60µm). **E-Eii** Hatchling postlarva 166dpf with mineralised dissoconch (DI) secreted by one cell thick folded mantle (Ei-Eii), siphon (Si), adductor muscle attachment (Ad) and digestive gland/stomach (Dig/sto) are visible through the transparent shell, foot (fo) active (scale bar 100µm). **F-Fii** Adult mantle with outer folds and fused inner fold, periostracal grooves secreting two-layered periostracum, scale bar = 2mm, reproduced with permission from Sleight et al. (2016a). **G** Schematised overview of mantle morphogenesis, mantle tissue black, periostracum grey. At late D-larvae stage PII (66dpf) mineralisation coincides with formation of folds in larval mantle edge. **H-Hii** SEM of late D-larval shell 66dpf showing delineated PI and PII shells, polygonal mottling on the dorsal surface suggestive of the calcification nucleation points, Hi & Hii show the shell is brittle and hence mineralised, H scale bar 100µm, Hi scale bar 35µm, Hii scale bar 10µm. **I-Iii** SEM of postlarval shell 160dpf DI delineated from PI and PII shells, arenophilic secretions and intraperiostracal spikes present. I scale bar 100µm, Ii scale bar 5µm, Iii scale bar 3µm. **J-Jii** SEM cross section of adult shell microstructure, a relatively think periostracum (J), an outer layer of aragonitic granular prisms (Ji) and an inner layer of aragonitic nacreous sheets (Jii), scale bars 5µm.

### Transcriptional characterisation of shell development in *Laternula elliptica*

Frist, a candidate gene approach was used to ask if the larval shell is built using the same protein coding genes as the adult shell. The expression of six genes encoding upstream regulatory proteins such as transcription factors and signalling molecules, and five downstream effector genes encoding extracellular matrix proteins and biomineralisation enzymes, were quantified over developmental time. All candidates had previously been characterised in adult shell secretion and repair in this species (via transcriptomics, computational gene network predictions, shell proteomics, qPCR and mRNA *in situ* hybridisations (Sleight et al. 2020; Sleight et al. 2016a; Sleight et al. 2015)). All of the upstream regulatory candidates were expressed during development, either increasing over time, or oscillating between stages (Fig. 2Bi). Only one of the downstream candidates, *le-meg*, was expressed during early shell development and the remaining four (*mytilin, pif, tyrA* and *tyrB*) were only expressed in the latest postlarva stage (Fig. 2Bii).

**Figure 2.**
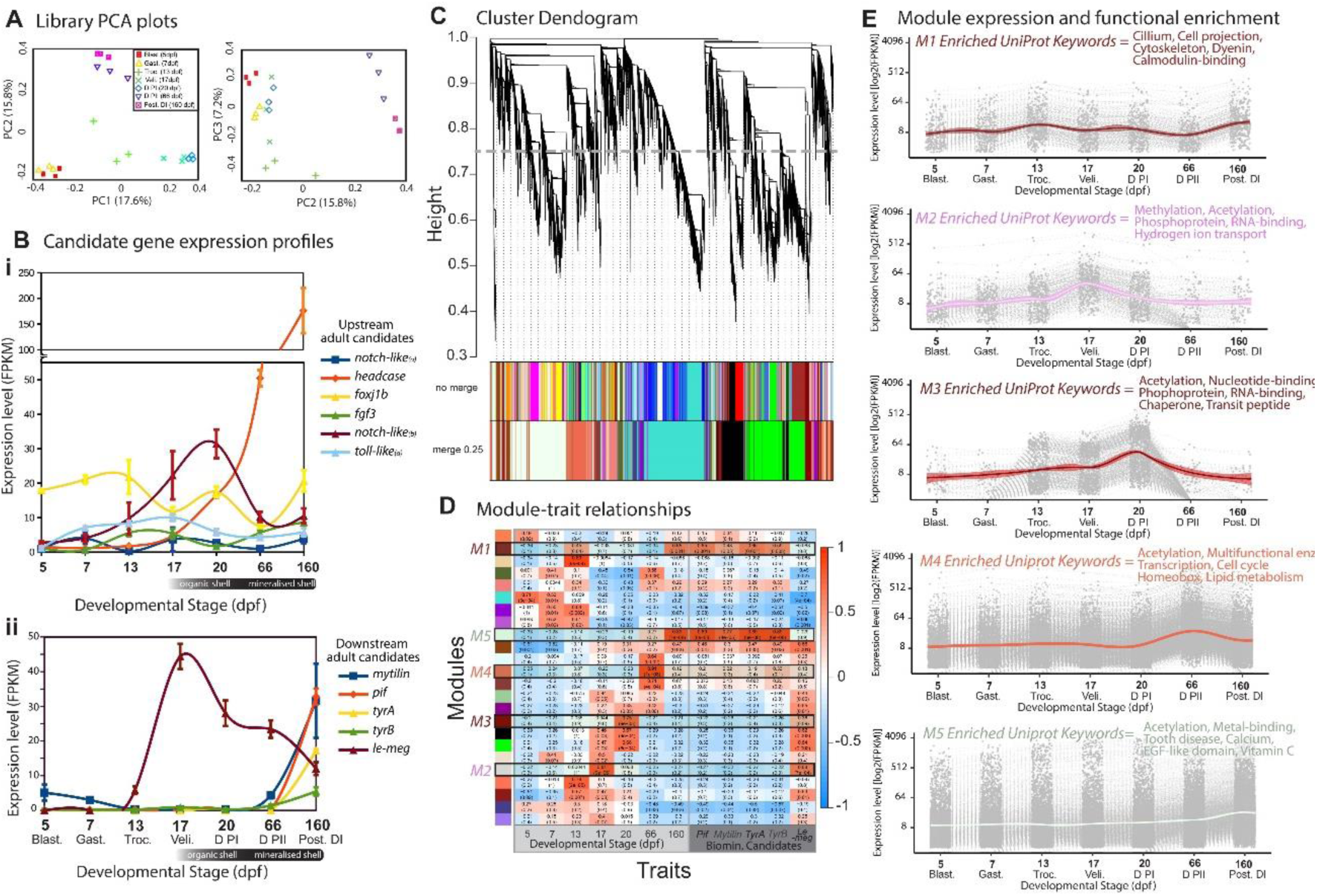
Candidate gene and trait-based approach to decipher transcriptional regulation of mantle and shell development in *Laternula elliptica* using bulk RNA-Seq. **A** PCA plots (PC1-3) of each RNA-seq library clustering by developmental stage (n=3 libraries per stage). **B** Average expression (Fragments Per Kilobase of transcript per Million [FPKM], mean average, +/-S.E., n=3) at each stage of shell development of candidate upstream regulatory and downstream effector genes. **C** Dendrogram obtained by average linkage hierarchical clustering using the Weighted Correlation Network Analysis (WGCNA) R package, module assignment determined by the Dynamic Tree Cut after merging at 75% cut-off. **D** Correlation of eigengene modules to shell development traits (dpf and expression of candidate downstream adult biomineralisation genes). Each row corresponds to an eigengene module and each column a trait, color-coded by direction and degree of the correlation (Pearson R, red = positive correlation; blue = negative correlation). Five eigengene modules are highlighted (M1-M5) as they significantly positively correlate to shell development traits of interest. **E** All transcripts in each eigengene module of interest (M1-M5) extracted and mean average FPKM plotted over developmental time (n=3, shaded +/-95% confidence interval, method = loess). Selected biologically interesting significantly enriched Uniprot keywords highlighted (calculated using String-DB, supplementary file 1).

Next, genes with developmental expression profiles indicating likely involvement in regulating shell development were screened. A trait-based approach, using Weighted Correlation Network Analysis (WGCNA) was used to cluster expression profiles into modules (Fig. 2C). Module-trait relationships were then calculated to identify sets of genes (eigengenes) whose expression significantly correlated to the previously identified mantle and shell development landmarks. Five eigengene modules (M1-5) were extracted based on their significant positive correlation to traits of interest (P<0.001, R>0.65, Fig. 2D-E). M1 contained 352 genes and was significantly positively correlated to the postlarva stage (DI, mineralised, intraperiostracal spikes and arenophillic secretions) and the average expression profile of two downstream adult candidate shell genes (*pif* and *tyrA*). M1 was enriched in genes related to the cytoskeleton, protein transport (dynein) and calmodulin-binding (conserved developmental genes in this module include: *arm, ltbp4, foxj1, vwa3a, iqcg, camk4*). M2 contained 337 genes whose expression profiles significantly positively correlated to veliger stage 17dpf (first organic PI initially deposited) and the average expression profile of one downstream adult candidate gene (*le-meg*). M2 was enriched in genes related to methylation and hydrogen ion transport (conserved developmental genes in this module include: *alpl, chs2, sbp1, h2b*). M3 contained 647 genes and was significantly positively correlated to early D-larvae (PI) stage 20dpf (first organic shell) but none of the adult candidate shell genes. M3 was enriched in genes related to protein folding and protein transport (conserved developmental genes in this module include: *hsp70, cpn601, hsp90, btf3, ccb23*). M4 contained 2,875 genes whose expression significantly positively correlated to late D-larvae (PII) stage 66dpf (mineralised shell) but none of the adult candidate shell genes. M4 was enriched in genes related to proliferation/growth, hox code transcription factors and multi-functional enzymes (conserved developmental genes in this module include: *a-somp, fgf1, abd-a, abd-b, nkx2*.*6, mox1*). M5 contained 6,369 genes significantly positively correlated to postlarva stage (DI) 160dpf and the average expression profile of four of the adult biomineralisation candidate genes (*pif, mytilin, tyrA* and *tyrB*). M5 was enriched in genes related to calcification, egf-like domains and vitamin C (conserved developmental genes in this module include: *bmp2, bmp3, bmp7, fgfl-1, ca, fz4, notch1, wnt4, wnt2b, tyr2, perl*). Modules 2-4 contain genes whose expression dynamics suggest they are involved in the regulation of early larval shell, where as M1 and M5 are both correlated to the downstream adult shell genes and show an increase in expression only in the latest stage, the postlarva (Supplementary file 1).

A second unbiased statistical approach was used to identify genes that were upregulated at each consecutive stage of shell development. Significantly upregulated genes were screened for functional categories relating to shell secretion (Fig. 3), extracted, and plotted over time in clustered heatplots (Fig. S1). Genes upregulated at the D-larvae (PII) stage included a striking number of transcription factors and signalling molecules, such as components of the hh and wnt signalling pathways (*gli, ptch, fzd4* and *wnt6*). The stage that had the fewest upregulated genes was D-larvae (PI) with 25 genes upregulated including genes involved in signalling (*rhoj* and *apolpp*), ion transport (*orct*) and the extracellular shell matrix (*pif*) (Supplementary file 1). Many of the screened biomineralisation genes had a stage-specific expression pattern. For example, genes that were highly upregulated in the postlarva stage (when the downstream adult shell candidates are also highly expressed) were expressed at very low levels through all of the earlier stages of shell development. Some of the stage-specific expression patterns were explained by ontogenetic partitioning of isoforms and/or paralogues (indicated with arrows Fig. S1), for example we found an isoform of *ptch3* that was expressed only in early shell development (significantly upregulated in the veliger stage) and a different isoform only expressed late in shell development (significantly upregulated in D-larvae PII stage).

**Figure 3.**
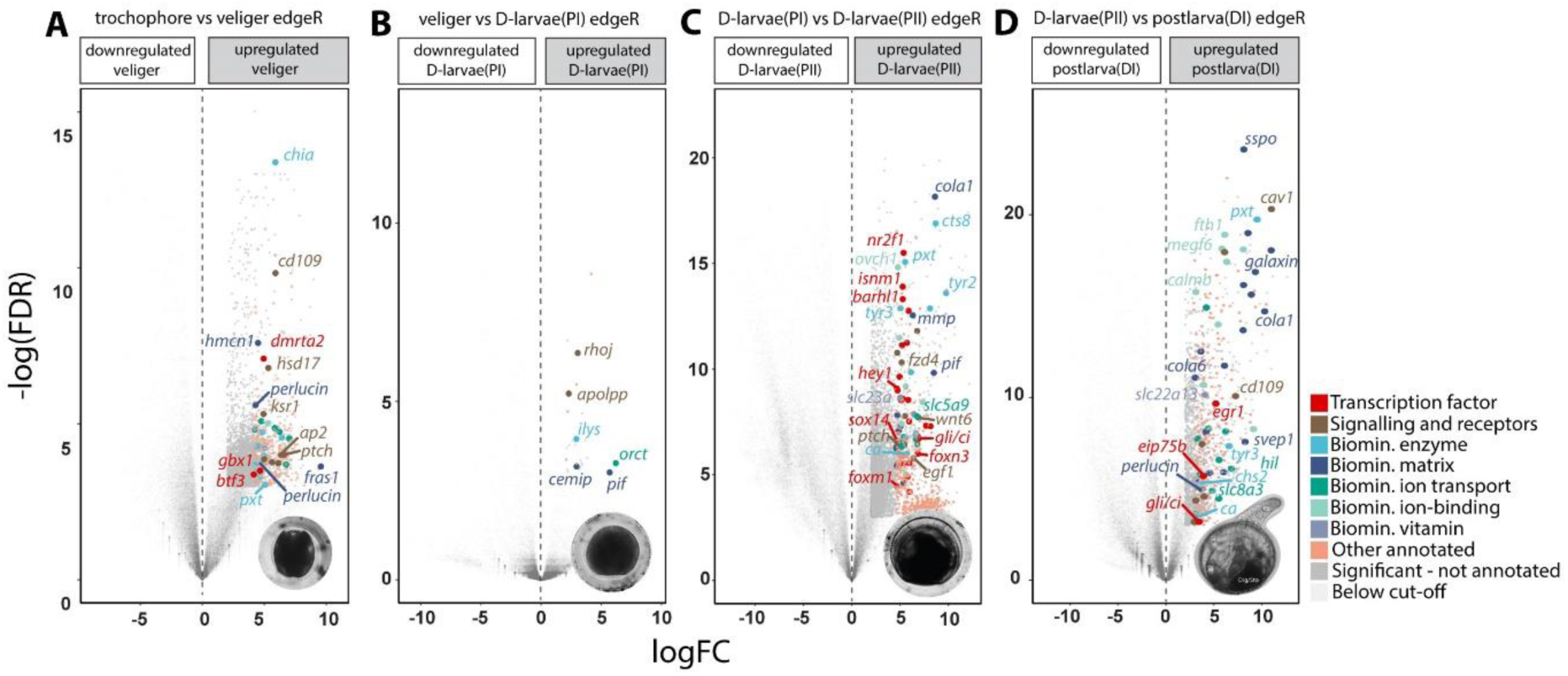
Unbiased pairwise differential expression approach to decipher transcriptional regulation of shell development in *Laternula elliptica*. Differential expression was determined using edgeR with a negative binomial additive general linear model and quasi-likelihood F-test, corrected for multiple testing using the Benjamin-Hochberg method to control the false discovery rate (FDR). Upregulated genes at each stage screened for statistical significance (FDR<0.05), magnitude (log_2_FC>2) and functional categories related to shell development, as per colour-coded key. Specific genes of biological interest are labelled. **A** upregulated genes at initial organic shell stage, veliger 17dpf. **B** upregulated genes at early D-larvae (PI) stage 20dpf with full organic PI shell. **C** upregulated genes at late D-larvae (PII) stage 66dpf when larval shell first begins to be mineralised and folds established in the larval mantle. **D** upregulated genes in hatchling postlarva (DI) stage 160dpf.

The signalling components upregulated during the shell development stages include hh and wnt pathways (Fig. 4A-B). To further explore the possible role of these signalling pathways in shell development we extracted genes relating to the canonical transduction of the hh and wnt pathways and plotted them over developmental time (Fig. S2). Most of the hh and wnt signalling genes were expressed prior to the secretion of the larval shell, followed by very low levels during early shell development stages– veliger and D larvae (PI) stages. One exception was a *ptch* isoform, which was upregulated again at the late D (PII) stage, coinciding with the beginning of larval mantle folding and onset of mineralisation.

**Figure 4.**
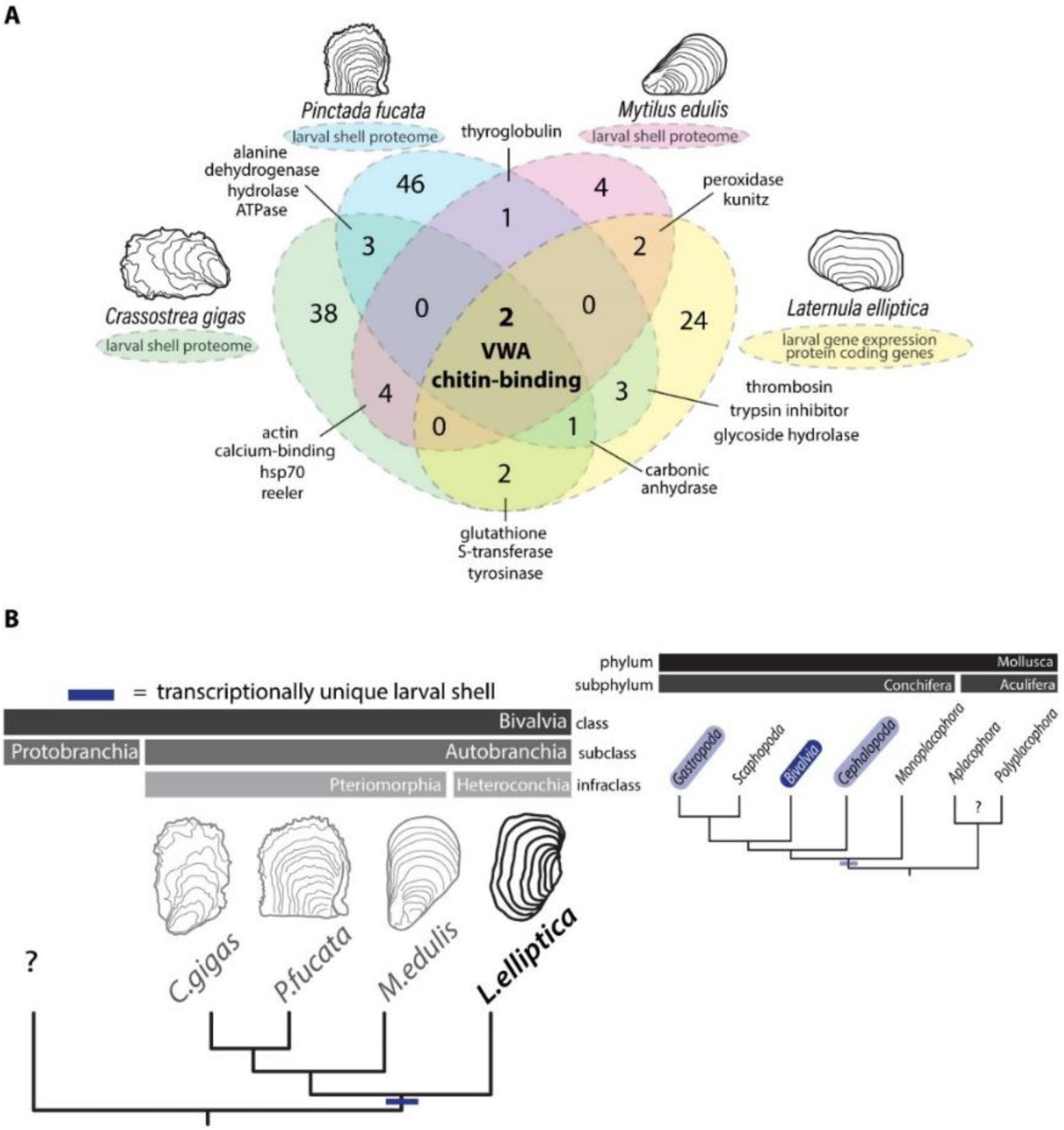
Development-specific downstream effectors control larval shell production in autobranchians, and more broadly across conchiferans. **A** Larval downstream effectors, such as shell matrix proteins and biomineralisation enzymes translated to amino acid sequences and surveyed for known domains. Protein domains compared against larval shell proteomes to search for conserved vs lineage specific domains using OrthoVenn2. **B** A development-specific larval shell proteome (versus adult shell proteome) in three species of Pteriomorhpia and, presented here, transcriptionally unique, development-specific downstream shell genes in a representative from Heteroconchia points to an ancestral condition to the Autobranchia subclass. Development-specific shell genes and isoforms have also been reported in gastropods. Taken together with many conchiferan molluscs retaining a larval shell but secondarily loosing, or reducing, an adult shell leads us to hypothesise that a transcriptionally unique larval shell is likely a deeply conserved trait to the Conchifera (no known developmental data for Aculifera or Protobranchia).

### Interspecies comparison of the molecular control of larval shell development

To test if larval shell matrix proteins are more conserved than those of adults, genes identified as likely larval shell downstream effectors in *L. elliptica* were compared with the three publicly available larval shell proteomes (*Crassostrea gigas, Pinctada fucata* and *Mytilus edulis*). Protein domains were compared across the four species to search for conserved vs lineage specific domains, as per Arivalagan et al. (2017). Only two protein domains were present in all four species larval shell datasets, VWA and chitin-binding, carbonic anhydrase was present in three out of four species (Fig. 4A).

## Discussion

We have quantified the ontogenetic expression dynamics of genes putatively involved in shell development for the first time in a heteroconchian bivalve, the Antarctic clam *L. elliptica*. These genes can be broadly categorised into two groups: downstream effectors that code for proteins involved in the extracellular shell matrix, enzymes and ion transporters (such as peroxidases, chitinases, matrix metalloproteinases, galaxins, ion and vitamin transporters) and upstream regulatory genes that code for transcription factors and signalling components (wnt and hh signalling components, hox, fox and sox transcription factors). Owing to a lack of genetically tractable model systems to conduct gene function experiments in molluscs (and more broadly spiralians), data on the gene regulatory networks that drive processes such as shell development and biomineralisation in this group are sparse, either focussing on two or three candidates (Kin et al. 2009; Liu et al. 2017; 2020; Nederbragt et al. 2002; Samadi and Steiner 2009; Tan et al. 2018), or use only computational methods (Sleight et al. 2020). Previous work has implicated bmp signalling in the regulation of molluscan shell secretion using *in situ* expression data and perturbations such as RNAi, DMH1 (a bmp inhibitor) or addition of bmp protein. Results have been variable; work in bivalves suggest bmp signalling could play a role in controlling adult shell secretion (Zhao et al. 2016) and shell development (Liu et al. 2020; Nederbragt et al. 2002). More recent experiments in the gastropod *Crepidula fornicata* found that when bmp signalling was disrupted during development, there was no observed effect on shell phenotype (Lyons et al. 2020) and single-cell RNAseq data from the trochophore stage of the heteroconchian bivalve *Dreissena rostriformis* suggests no bmp gene expression in shell-field cells (Salamanca-Díaz et al. 2022). The WGCNA analysis presented here found bmp pathway components were correlated only to the postlarva stage and the upregulation of the downstream adult shell candidates, but no bmp components were found using the differential expression approach. Taken together with previous reports in other groups, we hypothesise that bmp signalling is not involved in early shell development in heteroconchian bivalves, but could play a role in controlling the growth and maintenance of shell secretion at later stages.

The transcriptomic profiling data presented here suggests that hh and wnt signalling pathways are involved in shell development. These signalling pathways have well characterised roles in animal dorsal-ventral (wnt) and anterior-posterior patterning (hh), and the outgrowth of appendages in vertebrates and arthropods, as well as osteoblast differentiation and digit patterning in vertebrate endoskeletons (Hiscock et al. 2017; Riddle et al. 1993; Witte et al. 2009). Previous work has linked wnt signalling genes to biomineralisation in bivalves (Gao et al. 2016), and most recently, wnt and hh signalling have been shown to pattern cephalopod arm development (Tarazona et al. 2019). If cephalopod “limbs” evolved by parallel activation of a gene regulatory network for appendage development that was present in the bilaterian common ancestor (Tarazona et al. 2019), our data also suggest that this genetic program for appendage development may have been co-opted for shell development. More data is required on the spatial expression and function of hh and wnt genes in relation to mollusc shell development, and future work in species more suited to such methods should prioritise these studies.

Recent larval shell proteomes of three pteriomorphian bivalve species report that a development-specific repertoire of downstream effector proteins are found in the extracellular matrix of the larval shell when compared to the adult shell (Carini et al. 2019; Zhao et al. 2018). Here, we asked if this pattern holds true in a heteroconchian representative, and aimed to understand the transcriptional regulation of larval shell deposition versus that of adults. We find that previously characterised downstream adult shell genes are largely not involved in larval shell secretion (except for *le-meg*), neither the first organic stage nor early mineralised stages, but that the upstream regulatory genes, such as signalling molecules and transcription factors, are more conserved between life history stages. Our data suggest that global transcriptional mechanisms underpinning the unique larval shell proteins include ontogenetic partitioning of isoforms and paralogues, in addition to some *bona fide* unique larval genes (such as *btf3, pap18, chid1, cemip* and *nsf*). We hypothesise that, similar to other developmental systems, such as *svb*-driven gene regulatory networks in the formation of diverse actin rich projections in arthropods (Smith et al. 2018), conserved gene regulatory networks drive shell deposition in larval and adult life history stages, but the downstream effectors activated by the network depends on local factors in different life history and cellular contexts. As is the case with many systems studying morphological evolution, questions regarding the connection between gene regulatory networks and downstream effectors, the generation of temporal and local contexts and how different downstream effectors exert influences on phenotypic outcomes, remain unanswered.

Terminal downstream effectors in adult shells, such as extracellular shell matrix proteins, are rapidly evolving, lineage-specific and surprisingly diverse, with just four domains critical for all molluscan biomineralisation (carbonic anhydrase, chitin-binding, VWA and tyrosinase (Arivalagan et al. 2017)). To test if this pattern holds true for early life history stages where, phenotypically at least, shells are more conserved between species, we carried out comparative domain analysis between heterochoncian and pteriomorphian bivalves. Similar to the pattern in adult life history stages for these groups, we find only two domains shared between the four species (chitin-binding and VWA domains), despite the similarity in shell phenotype across species at this stage. In our analysis we compared data generated by different groups using different methods, in addition, our data is a prediction of biomineralisation effectors and hence, it is likely that the number of conserved domains are underestimated. For example, it is unlikely that carbonic anhydrase is truly absent in larval mussel shells as reported Carini et al., (2019), but rather it was technologically difficult to detect all proteins present in larval shell with the small input material that is available from developmental stages.

For the first time, we have compared the transcriptional regulation of shell secretion at different life history stages of a heteroconchian bivalve. Studying a heterconchian representative, and comparing it to data available for pteromorphians allows us to hypothesise that a development-specific transcriptionally unique larval shell is likely an ancestral feature of autobranchians, or perhaps even more broadly to conchiferans (Fig. 5B). Ontogenetic partitioning of some specific shell gene isoforms has been reported in gastropods, as well as alternative splicing to generate diverse shell matrix proteins from a single genomic loci (*Halitosis asinine* and *Lymnaea stagnalis* (Herlitze et al. 2018; Jackson et al. 2006)). In addition, some conchiferan molluscs have a larval shell but have secondarily lost, or reduced, an adult shell (Knutson et al. 2020). There is a huge diversity of form and function between different life history stages in molluscs. Taken together, these findings lead us to hypothesise that a development-specific transcriptionally unique larval shell may be a deeply conserved trait to the Conchifera, but more data from diverse taxa, especially the Acuilifera, are needed to resolve questions on the evolution of biomineralisation in the molluscs.

## Methods

### Animal husbandry, spawning and developmental staging

Embryos were obtained from an adult broodstock of sexually mature *L. elliptica* individuals and divided into three independent closed-system 1L tanks. Embryos were maintained at 0°C ± 0.5°C, aerated with an airstone with 50% water changes every two days using autoclaved seawater. Embryos were staged as per Peck et al. (2007).

### Fixation, histology and imaging

For histology and Scanning Electron Microscopy (SEM), embryos were fixed in 500 µL of 2.5% glutaraldehyde in phosphate buffered saline (PBS) at room temperature for 30 minutes, rinsed twice in PBS with 0.1% tween, dehydrated into 100% ethanol and stored at 4°C.

For histology, embryos were cleared in Histosol (National Diagnostics, 3 × 20 min, room temperature), transitioned through 1:1 Histosol:molten paraffin (2 × 30 mins, 60°C) then pure molten paraffin (RA Lamb Wax – Fisher Scientific, 60°C overnight). Five changes of molten paraffin (each >1 hr) were conducted before embedding in a Peel-A-Way embedding mold (Sigma). Wax blocks were serially sectioned at 8 μm on a Leica RM2125 rotary microtome. Sections were stained with a modified Masson’s trichrome stain as per Witten and Hall (2003). All histology was conducted on a minimum of 3 individuals per stage. Sections were imaged on a Zeiss Axioscope A1 with Zen software (at the University of Cambridge).

For SEM of the shell late D (66dpf) and postlarval (160dpf) stages, larvae were transferred from ethanol to electron microscope stubs, attached using carbon adhesive discs, carbon coated and examined using a QEMSCAN 650F (at the University of Cambridge) at accelerating voltages of 10 kV.

Each stage was also live imaged on an upright compound light microscope fitted with a camera (Olympus BX50 microscope fitted with Olympus PM-C35 camera using Olympus U-CMAD-2 software, at the British Antarctic Survey) by mounting in seawater onto a glass slide, under a coverslip on small clay feet. All live imaging was conducted on a minimum of 3 individuals per stage.

Images were adjusted for contrast and brightness, cropped, rotated, and flipped. All plates were constructed in Adobe Illustrator software.

### Sampling, RNA extraction and sequencing

Triplicate RNA-Seq samples were collected for each developmental stage, one from each independent tank. For each sample, exactly two hundred staged-matched embryos were snap frozen in a 70% ethanol dry ice slurry and stored at -80°C. Total RNA was extracted from each sample as per manufactures recommendations (Relia Miniprep kit, Promega) and tested for quality and quantity using Nanodrop and Agilent Tapestation. All samples had an RNA Integrity Number (RIN) of over 7. Libraries were prepared by the DNA sequencing facility in the Biochemistry Department at the University of Cambridge (TruSeq Stranded mRNA, Illumina) and sequenced on an Illumina NextSeq500 generating over 300 million 150bp stranded paired-end reads. All raw data is publicly available from NCBI SRA accession number: PRJNA803976.

### Bioinformatics and statistical analyses

Raw reads (total 309,593,642) were cleaned using ea-utils tool v1.1.2 fastq-mcf (quality –q 30, and length –l 100), after cleaning 296,480,254 reads remained. Prior to *de novo* assembly, reads were normalised using the Trinity v.2.2.0 utility script (insilico_read_normalization.pl – max_cov 50), leaving 64,355,068 reads for assembly. Clean, normalised reads were assembled using Trinity v.2.2.0 with default parameters (Grabherr et al. 2011; Haas et al. 2013). Assembly quality was assessed using Trinity utilities and the gVolante webserver tool (Table S1). Transcript abundance was estimated by alignment-based quantification using Trinity v.2.2.0 utilities. Transcripts from each cleaned library were aligned to the transcriptome using bowtie2 with default parameters and transcript abundance estimates were calculated using RNA-Seq by Expectation-Maximization (RSEM). Raw counts and Trimmed Mean of M-values [TMM] normalised Fragments Per Kilobase Of Exon Per Million Fragments Mapped [FPKM] matrices were generated.

Weighted correlation network analysis (WGCNA) was used to find clusters, termed modules, of genes with highly correlated expression across all libraries. Each cluster was then correlated to external traits of interest (Langfelder and Horvath 2008). The raw counts matrix was loaded in R and EdgeR (Robinson et al. 2010) functions were used to remove lowly expressed transcripts (keep transcripts with cpm >5 in ≥4 libraries) leaving 37,000 transcripts for WGCNA. TMM-FPKM values were used to calculate a gene dissimilarity matrix (adjacency= softpower 16 and signed, TOMsimilarity = signed) and hierarchical clustering was performed (method = average). Modules were determined using the cutreeDynamic function with a minimum gene membership threshold of 30 and dynamic tree cut-off of 25. Modules were correlated to external traits (days post fertilization or average candidate gene expression values). Modules of that were significantly correlated to traits of interest were extracted and all transcripts were putatively annotated based on sequence similarity searched using blastx against Uniprot (http://www.uniprot.org/), and tested for functional enrichment.

Pairwise differential gene expression tests were performed to find transcripts that were upregulated at each stage of shell development (compared to the previous stage). Using the EdgeR package, a negative binomial additive general linear model with a quasi-likelihood F-test was performed and p-values were adjusted for multiple testing using the Benjamini-Hochberg method to control the false discovery rate, cut-offs for statistical significance (FDR ≤0.05) and magnitude were used (log_2_FC>2). Upregulated transcripts were putatively annotated based on sequence similarity searched using blastx against Uniprot (http://www.uniprot.org/), and screened for functional categories relating to gene regulation and shell secretion.

All upregulated genes that were functionally categorised as downstream effectors, such as shell matrix proteins and biomineralisation enzymes were translated to amino acid sequences and surveyed for known domains. Protein domains were then compared against published larval shell proteomes for oyster (Zhao et al. 2018) and mussel (Carini et al. 2019) species to search for conserved vs lineage specific domains as per Arivalagan et al., (2017), using OrthoVenn2.

## Data Availability

Raw read data generated in this publication is freely available at NCBI SRA with the following accession PRJNA803976. All analysis scripts and results are available in Supplementary File 1.

## Acknowledgements and Funding Information

We are grateful to Andrew Gillis for use for paraffin histology and microscopy equipment and enthusiastic support of this work and the Rothera marine team for collecting the adult *Laternula elliptica* broodstock.

This work was supported by UKRI Natural Environment Research Council (NERC) Core Funding to the British Antarctic Survey, a DTG Studentship (Project Reference: NE/J500173/1) and a Junior Research Fellowship to VAS from Wolfson College, University of Cambridge.

## Author Contributions

**AC** Performed all wet-lab experiments, contributed to imaging, data analysis and interpretation, and reviewed and edited the manuscript.

**MSC** Contributed to supervision of AC, funded the sequencing, contributed to wet-lab experiments, imaging and data interpretation, reviewed and edited the manuscript.

**EH** Conducted all SEM imaging, contributed to data interpretation and reviewed and edited the manuscript.

**LSP** Conducted adult broodstock collection, contributed to wet-lab experiments and data interpretation, reviewed and edited the manuscript.

**VAS** Conceptualized and oversaw the study, supervised AC, contributed to wet-lab experiments and imaging, conducted all data analysis and interpretation, prepared all figures and wrote the manuscript.

## Supplementary Figures, Tables and Files

**Figure S1.**
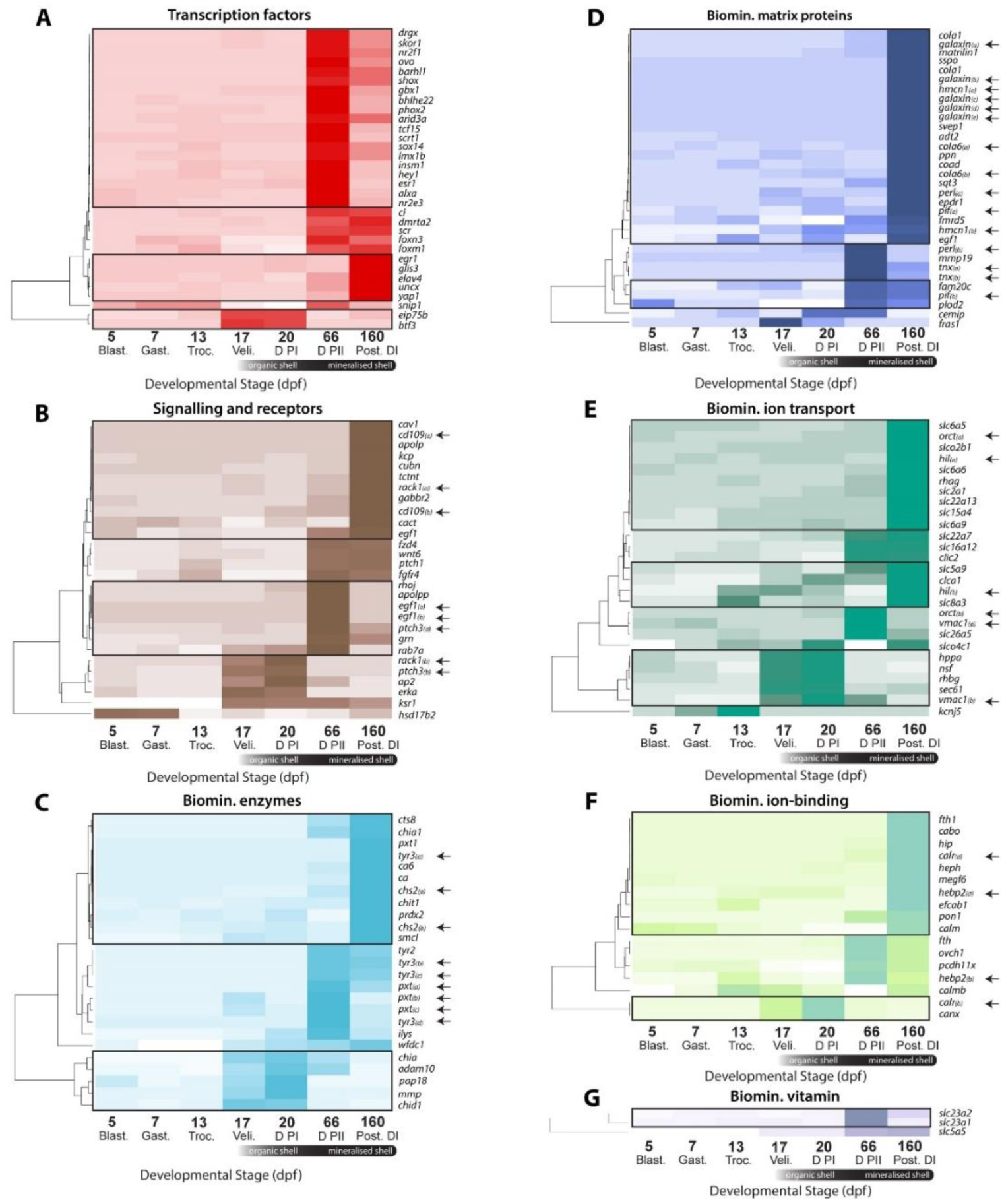
Clustered heatplots of significant functionally screened genes from unbiased differential expression analysis, genes typically show stage-specific expression patterns. Mean average FPKM (n=3) z-scaled over developmental time and clustered by temporal expression profile on y axis only using hclust (method = single, clustering represented by dendograms on left). Multiple isoforms of the same gene with unconfirmed nomenclature labelled with lowercase letters in parentheses and highlighted with arrows.

**Figure S2.**
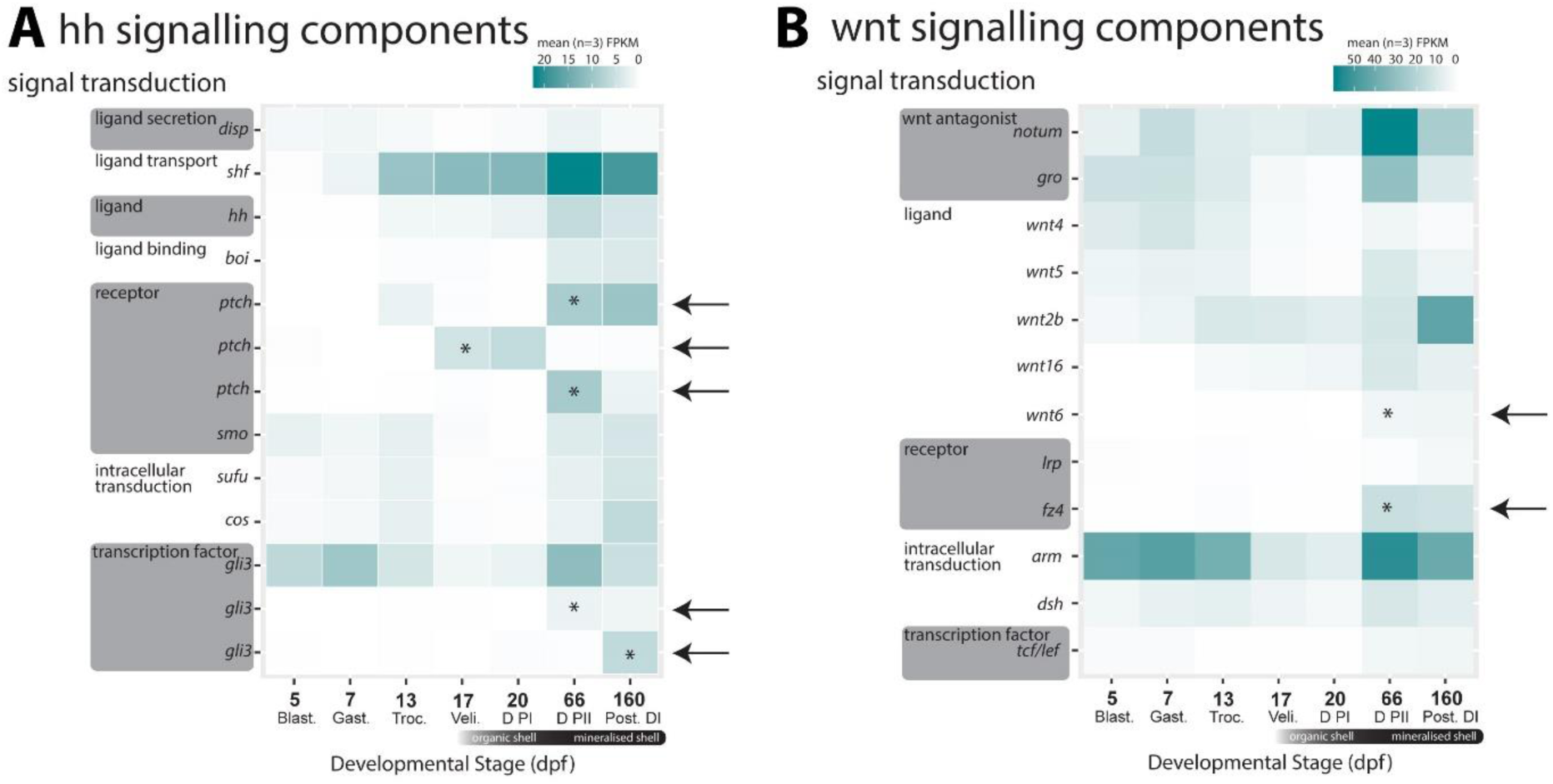
Both hh and wnt signalling pathway genes were significantly upregulated during shell development and could be involved in patterning and proliferation of the larval shell field and larval mantle. A Heatplot of hh signalling genes and B wnt signalling genes. Statistically significantly upregulated genes (edgeR, FDR<0.05, log_2_FC>2) identified with arrows and stage at which significantly upregulated marked with an asterisks (*). Order of genes on the y axis presented in terms of signal transduction cascades from ligand secretion towards the top and recruitment of downstream transcription factors to the nucleus at the bottom.

**Table S1.** Summary statistics for *de novo* assembled *Laternula elliptica* developmental transcriptome.

| **Reads:** |  |
| --- | --- |
| Raw reads | 309,593,642 |
| Clean reads | 296,480,254 |
| Normalised reads | 64,355,068 |
| **Assembly:** |  |
| Total trinity transcripts | 964,764 |
| Total trinity genes | 666,835 |
| % GC | 46.3 |
| **Statistics based on all isoforms per gene:** |  |
| N50 | 720 |
| Median length | 462 |
| Mean average length | 662 |
| **Read representation (average read content per library):** | 67% |
| **BUSCO v3, Metazoa Gene Set (n=978)** | C:99.1%[S:30%,D:69%] F:0.7%,M:0.2% |

**Supplementary file 1**. Data archive including readme.txt file detailing curation, includes annotations, statistical analysis R scripts and results tables.

