## Supplementary File 1 for "Evolutionary conservation and divergence of the transcriptional regulation of bivalve shell secretion across life history stages": edgeR_script_pairwise_comparisons_DGE.R

```
#start by loading all the different libraries needed for statistical testing and plotting

library(sva)
library(edgeR)
library(ggplot2)
library(statmod)
library(dict)
library(naniar)
library(data.table)
library(heatmap.plus)
library(RColorBrewer)
library(reshape2)
library(wesanderson)
library(ggsci)
library(ggrepel)
library(systemfonts)
library(svglite)

#READ IN THE DATA - A FILE IN THE CURRENT WORKING DIRECTORY, IT IS RAW GENE COUNTS CALCULATED BY RSEM 
latty_bros_data <-read.delim("Raw_gene_counts.matrix", check.names=FALSE, stringsAsFactors=FALSE)

#MAKE DGE LIST AND TELL IT THE NAMES OF THE COLLUMNS
y <- DGEList(counts=latty_bros_data[,2:22], genes=latty_bros_data[,1])

#Add grouping, manually instruct which collum is which sample
group <- c("Blastula","Gastrula","Early_Trocophore","Veliger","Dlarvae","Late_Dlarvae","Blastula","Gastrula","Early_Trocophore","Veliger","Dlarvae","Late_Dlarvae","Blastula","Early_Trocophore","Veliger","Dlarvae","Late_Dlarvae","Gastrula","Postlarva","Postlarva","Postlarva")
y <- DGEList(counts=latty_bros_data[,2:22], genes=latty_bros_data[,1], group=group)

#NORMALISE THE DATA
y$samples$lib.size <- colSums(y$counts)
y <- calcNormFactors(y)
y$samples

#FILTER THE DATA TO REMOVE GENES WITH VERY VERY LOW COUNTS IN A THIRD OF THE LIBARIES ie must have over 0.1 cpm in 6 or more libaries 
keep <- rowSums(cpm(y)>0.1) >=6
y <- y[keep, , keep.lib.sizes=FALSE]

#THEN FILTER THE DATA TO REMOVE GENES WITH QUITE LOW COUNTS IN SOME LIBRARIES ie must have over 5 cpm in 3 or more librares
keep <- rowSums(cpm(y)>5) >=3
y <- y[keep, , keep.lib.sizes=FALSE]

#RE-NORMALISE FOR NEW FILTERED LIBRARY (PROBABLY NEGLIGIBLE BUT GOOD PRACTISE)
y$samples$lib.size <- colSums(y$counts)
y <- calcNormFactors(y)

#How any genes do I have left after this filtering?
summary(y$genes)

#DEFINE THE MODEL DESIGN
design <- model.matrix(~0+group, data=y$samples)
colnames(design) <- levels(y$samples$group)
design

#ESTIMATE DISPERSION
y <- estimateDisp(y,design, robust=TRUE)

#PLOT DISPERSION
tiff("Latty_bros_filtered_Robust_BCV.tiff")
plotBCV(y)
dev.off()

### first fit negative binomial GLM for each tag and produces an object of class DGEGLM (some new components)

fit <- glmQLFit(y, design)


###pass DGEGLM object to glmQLFTest() to carry out the desired QL F-test.
Troc_vs_Veliger <- makeContrasts(Veliger-Early_Trocophore, levels=design)

Vel_vs_D <- makeContrasts(Dlarvae-Veliger, levels=design)

Dlarvae_vs_LateDlarvae <- makeContrasts(Late_Dlarvae-Dlarvae, levels=design)

LateD_vs_postlarva <- makeContrasts(Postlarva-Late_Dlarvae, levels=design)

########################

troc_vel_qlf <- glmQLFTest(fit, contrast=Troc_vs_Veliger)

Vel_D_qlf <- glmQLFTest(fit, contrast=Vel_vs_D)

D_vs_lateD_qlf <- glmQLFTest(fit, contrast=Dlarvae_vs_LateDlarvae)
  
LateD_postlarva_qlf <- glmQLFTest(fit, contrast=LateD_vs_postlarva)

######################

tt_troc_vel_qlf <- topTags(troc_vel_qlf, n=NULL)

tt_Vel_D_qlf <- topTags(Vel_D_qlf, n=NULL)

tt_D_vs_lateD_qlf <- topTags(D_vs_lateD_qlf, n=NULL)

tt_LateD_postlarva_qlf <- topTags(LateD_postlarva_qlf, n=NULL)

######################

    write.table(tt_troc_vel_qlf, file="tt_troc_vel_qlf.txt", row.names=FALSE, col.names=TRUE)
    
    write.table(tt_Vel_D_qlf, file="tt_Vel_D_qlf.txt", row.names=FALSE, col.names=TRUE)
    
    write.table(tt_D_vs_lateD_qlf, file="tt_D_vs_lateD_qlf.txt", row.names=FALSE, col.names=TRUE)
    
    write.table(tt_LateD_postlarva_qlf, file="tt_LateD_postlarva_qlf.txt", row.names=FALSE, col.names=TRUE)

######################

troc_vel_0.001 <- decideTestsDGE(troc_vel_qlf, adjust.method="BH", p.value=0.001)

Vel_D_0.001 <- decideTestsDGE(Vel_D_qlf, adjust.method="BH", p.value=0.001)

D_vs_lateD_0.001 <- decideTestsDGE(D_vs_lateD_qlf, adjust.method="BH", p.value=0.001)

LateD_postlarva_0.001 <- decideTestsDGE(LateD_postlarva_qlf, adjust.method="BH", p.value=0.001)

#####################

summary(troc_vel_0.001)

summary(Vel_D_0.001)

summary(D_vs_lateD_0.001)

summary(LateD_postlarva_0.001)

##################


sessionInfo()
```
