## Supplementary File 1 for "Evolutionary conservation and divergence of the transcriptional regulation of bivalve shell secretion across life history stages": README.txt

This data archive includes files relating to the analysis of Laternula elliptica developmental transcriptomics data. There are two directories:

-WGCNA

-edgeR_DE

Below is a brief outline of the contents in each directory:


-WGCNA
In this directory there is a commented R script used for analysis.
There is also a further sub directory named "Modules"

	-Modules
	This directory includes two subdirectories:
		
		-Annotations
		This directory includes annotation (swissprot human) table for genes in each module

		-Enrichment_analysis_results
		A table for each module's enrichment results

-edgeR_DE
This directory includes the R script used to run the differential gene expression analysis "edgeR_script_pairwise_comparisons_DGE.R" and the compiled results in excel table format, one tab per pairwise comparison "edgeR_DGE_results_tabs.xlsx"
