## Supplementary File 1 for "Evolutionary conservation and divergence of the transcriptional regulation of bivalve shell secretion across life history stages": WGCNA_analysis_v2.1_merge_modules.R

```
# Load WGCNA, flashClust libraries, expression and trait data
library(WGCNA)
library(flashClust)
allowWGCNAThreads()

load(file="expressiontraitdata_feb13th2020.RData")

####################################################################################################
###############################Data now ready for analysis##########################################
####################################################################################################

#####################################################
# Choose a soft threshold power- USE A SUPERCOMPUTER#
#####################################################
##chosen 16 as more than 20 samples and signed network as per FAQs when can't get a good scale free topology R^2

############################################## 
# Construct Networks- USE A SUPERCOMPUTER IRL#
##############################################


#build a adjacency "correlation" matrix
enableWGCNAThreads()
softPower = 16
adjacency = adjacency(datExpr, power = softPower, type = "signed") #specify network type
head(adjacency)

#translate the adjacency into topological overlap matrix and calculate the corresponding dissimilarity:
TOM = TOMsimilarity(adjacency, TOMType="signed") # specify network type
dissTOM = 1-TOM

#################### 
# Generate Modules #
####################
 
# Generate a clustered gene tree

geneTree = flashClust(as.dist(dissTOM), method="average")
plot(geneTree, xlab="", sub="", main= "Gene Clustering on TOM-based dissimilarity", labels= FALSE, hang=0.04)
#This sets the minimum number of genes to cluster into a module
minModuleSize = 30
dynamicMods = cutreeDynamic(dendro= geneTree, distM= dissTOM, deepSplit=2, pamRespectsDendro= FALSE, minClusterSize = minModuleSize)
dynamicColors= labels2colors(dynamicMods)
MEList= moduleEigengenes(datExpr, colors= dynamicColors,softPower = softPower)
MEs= MEList$eigengenes
MEDiss= 1-cor(MEs, use = "pairwise.complete.obs")
METree= flashClust(as.dist(MEDiss), method= "average")
save(dynamicMods, MEList, MEs, MEDiss, METree, file= "Network_allSamples_signed_RLDfiltered.RData")
#save progress
  
#plot tree showing how the eigengenes cluster together
pdf(file="clusterwithoutmodulecolors.pdf")
plot(METree, main= "Clustering of module eigengenes", xlab= "", sub= "")
#set a threhold for merging modules at 0.25
MEDissThres = 0.25
merge = mergeCloseModules(datExpr, dynamicColors, cutHeight= MEDissThres, verbose =3)
mergedColors = merge$colors
mergedMEs = merge$newMEs
dev.off()
 
#plot dendrogram with module colors below it
pdf(file="cluster.pdf")
plotDendroAndColors(geneTree, cbind(dynamicColors, mergedColors), c("Dynamic Tree Cut", "Merged dynamic"), dendroLabels= FALSE, hang=0.03, addGuide= TRUE, guideHang=0.05)
moduleColors = mergedColors
colorOrder = c("grey", standardColors(50))
moduleLabels = match(moduleColors, colorOrder)-1
MEs = mergedMEs
dev.off()
#save progress 
save(MEs, moduleLabels, moduleColors, geneTree, file= "Network_allSamples_signed_RLDfiltered_merged.RData")


########################################
### Relate modules to external traits###
########################################

#Define number of genes and samples
nGenes = ncol(datExpr)
nSamples = nrow(datExpr)
#Recalculate MEs with color labels
MEs0 = moduleEigengenes(datExpr, moduleColors)$eigengenes
MEs = orderMEs(MEs0)
moduleTraitCor = cor(MEs, datTraits, use= "p")
moduleTraitPvalue = corPvalueStudent(moduleTraitCor, nSamples)
 
 
#Print correlation heatmap between modules and traits
textMatrix= paste(signif(moduleTraitCor, 2), "\n(",
                        signif(moduleTraitPvalue, 1), ")", sep= "")
dim(textMatrix)= dim(moduleTraitCor)
par(mar= c(6, 8.5, 3, 3))
 
 
#display the corelation values with a heatmap plot
pdf(file="heatmap.pdf")
labeledHeatmap(Matrix= moduleTraitCor,
            xLabels= names(datTraits),
            yLabels= names(MEs),
            ySymbols= names(MEs),
            colorLabels= FALSE,
            colors= blueWhiteRed(50),
            textMatrix= textMatrix,
            setStdMargins= FALSE,
            cex.text= 0.5,
            zlim= c(-1,1),
            main= paste("Module-trait relationships"))
dev.off()
#pull out genes belonging to a certain module, you can use the following command:
#names(datExpr)[moduleColors=="brown”]
```
